# The effect of feeding on microbiome and biogas composition in anaerobic CSTR

**DOI:** 10.1101/2025.02.07.637154

**Authors:** Georgios Samiotis, Manthos Panou, Vassiliki Tsioni, Themistoklis Sfetsas

## Abstract

his study investigates the performance of two Continuous Stirred Tank Reactors (CSTRs) with a focus on biogas yield, physicochemical parameters, and microbial dynamics. By integrating experimental observations with insights from recent literature, the research aims to elucidate the intricate relationships between reactor conditions, microbiome composition, and biogas production efficiency. Two substrates were used: a control substrate (SB1) and a protein-rich test substrate (SB2). The study monitored key parameters such as pH, Total Alkalinity of Carbonates (TAC), and volatile fatty acids (FOS), and analyzed the microbial communities using high-throughput sequencing. Results indicated significant temporal variations in pH, TAC, and nitrogen levels, with a declining FOS/TAC ratio. The introduction of SB2 led to increased biogas production and methane content, particularly at higher Hydraulic Retention Times (HRT). The study also highlighted the role of specific microbial taxa in enhancing biogas quality. These findings contribute to the development of optimized strategies for sustainable biogas production and process control.

## 1. Introduction

Biogas production has emerged as a pivotal technology in addressing global energy challenges while mitigating waste management issues. Anaerobic digestion (AD) is the cornerstone of biogas production, where organic matter undergoes microbial decomposition in the absence of oxygen, resulting in methane-rich biogas. Among various reactor configurations, the Continuous Stirred Tank Reactor (CSTR) has gained prominence due to its robust design, operational flexibility, and high efficiency in biogas generation. The CSTR provides continuous mixing of substrates and microbial populations, ensuring homogeneous conditions that optimize digestion rates and biogas yields (Nasir et al., 2012).

CSTRs are characterized by their ability to process a wide range of feedstocks, including agricultural residues, food waste, and livestock manure. This versatility, combined with their scalability and adaptability to varying operational parameters such as hydraulic retention time (HRT) and organic loading rate (OLR), makes CSTRs a preferred choice in industrial and research settings (Kaparaju et al., 2009). Notably, innovative configurations such as serially connected CSTR systems have demonstrated superior biogas yields by improving the utilization of volatile fatty acids (VFAs) and stabilizing the digestion process (Boe & Angelidaki, 2009).

Beyond reactor design and operational conditions, the composition and dynamics of the microbial community within the digester are critical to the success of the anaerobic digestion process. The synergistic interactions among hydrolytic, acidogenic, acetogenic, and methanogenic microbial populations determine the efficiency of substrate conversion into biogas. Advances in microbiome analysis have facilitated a deeper understanding of these microbial interactions, revealing strategies to enhance reactor performance. High-throughput sequencing techniques have identified specific microbial taxa associated with higher methane production, paving the way for targeted bioaugmentation and microbial management strategies (Parawira, 2012).

Microbial communities within CSTRs play a pivotal role in the anaerobic digestion process. Recent advances in microbial analysis, including high-throughput sequencing and metagenomics, have provided deeper insights into the composition and functionality of microbial consortia. For instance, studies have identified Clostridia and Methanobacteria as key contributors to the breakdown of complex organic substrates and methane production (Klocke et al., 2007). Such insights are critical for bioaugmentation and other strategies aimed at enhancing reactor performance.

Recent studies have also emphasized the importance of substrate characteristics and co-digestion strategies in boosting biogas yields. The inclusion of nutrient-rich co-substrates, such as livestock manure combined with industrial or agricultural residues, has demonstrated significant improvements in methane output and process stability (Romero & Romaní, 2024). Moreover, the use of artificial intelligence (AI) and modeling approaches in predicting and optimizing reactor performance has emerged as a transformative tool, allowing researchers to simulate different scenarios and enhance process efficiency (Fajobi et al., 2022).

This study explores the performance of two CSTR platforms with a focus on biogas yield, physicochemical parameters, and microbial dynamics. By integrating experimental observations with insights from recent literature, this research aims to elucidate the intricate relationships between reactor conditions, microbiome composition, and biogas production efficiency. The findings are expected to contribute to the development of optimized strategies for sustainable biogas production and process control.

## 2. Materials and Methods

### 2.1. Experimental design

In this study, two identical, 10 L volume, continuously stirred tank anaerobic bioreactors (CSTR1 & CSTR2) were operated in parallel. Both bioreactors were heated using a heating-blanket, and were thermostatically controlled at operational temperature of 37°C ± 0,5°C.

Mesophilic anaerobic biomass inoculums of 10 L, obtained from a 1MW biogas plant (Biogas Lagada SA) that is applying wet fermentation process, were used to fill each CSTR bioreactor. Feeding and discharging of the bioreactors was performed automatically (daily), and simultaneously, by individual peristaltic pumps set for CSTR1 and CSTR2.

Two feeding substrates were used, (i) a first substrate (SB1) obtained from the feeding mixture of Biogas Lagada SA, which was used during the start-up and system control period, when biomass acclimatization occurs, technical evaluation of the system is performed, and bioreactors benchmark conditions are obtained, (ii) and a second, protein-rich test substrate (SB2) composed from:

- Poultry manure from a broiler farm (poultry manure—PM).
- Swine manure from a pig farm (swine manure—SM).
- Cattle manure from a bovine farm (cattle manure—CM).
- Expired or unsuitable for human consumption food waste (FW), including animal-origin and plant-based liquids (e.g., milk, juices, oil), restaurant leftovers, fruits, vegetables, legumes, canned goods, deli meats, and dairy products The exact composition of SB1 and SB2 is presented in Table 1.

**Table 1.**
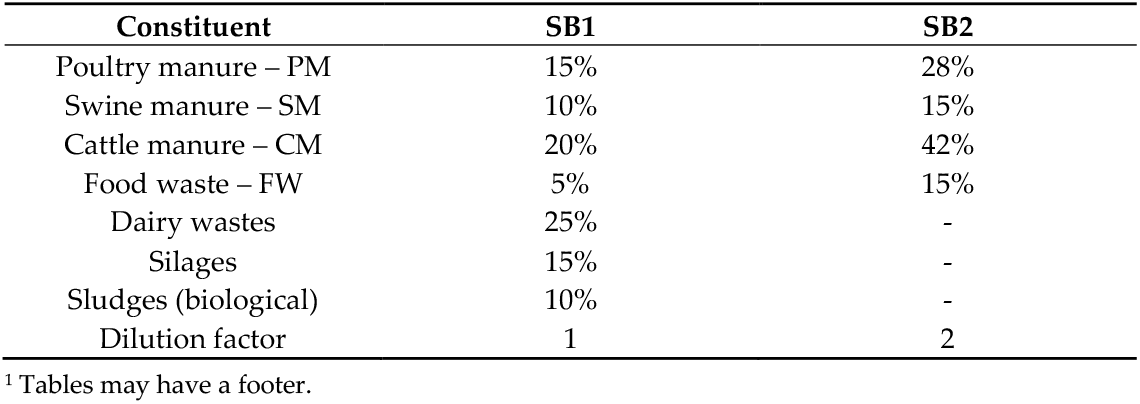
Composition of SB1 and SB2 feeding substrates.

Regarding SB2 preparation, after sampling, all raw materials were stored at −18°C, then grounded with a knife mill (Fritsch PULVERISETTE 11) and the use of dry ice to achieve a more homogeneous form. The applied process is in accordance with the requirements of DIN 4630/2016, which specifies standards for the homogenization and uniformity of samples intended for anaerobic digestion, as well as in compliance with EU Regulation 142/2011, which mandates that the particle size of raw materials for anaerobic digestion should not exceed 12 mm.

The duration of the experiment was 84 days, in the period which controlled changes were imposed on feedstock type, as well as on the operational hydraulic retention time (HRT). In more detail, during the first 24 days, SB1 was fed in CSTR1 and CSTR2 at a rate of 500 ml/d to maintain an HRT of 20 days. This is the exact HRT of the bioreactor of the biogas plant, where both inoculum and SB1 are obtained from. Thus, the technical evaluation of the system can be performed, and bioreactors benchmark conditions can be obtained, by comparing specific biogas production rates, as well as biogas composition, between the experimental setups, and the full-scale bioreactor (4000 m^3^).

After the initial 24-day experimental period, the second feedstock (SB2) was introduced on CSTR1 and CSTR2. HRT was maintained at 20 days for 18 days, and thereafter set to 30 days for a period of approximately three weeks (21 days) by lowering feeding rate to approximately 333 ml/d. Finally, a period of increasing HRT from 30 days to approximately 50 days was imposed during the last 21 days of the experiment by inducing starvation conditions, i.e., seize of feeding for another 21 days period.

### 2.2. Sampling and analysis

A plethora of physicochemical and microbiological parameters were analyzed in used feedstocks, and bioreactors’ material, in order to assess the impact of feedstock composition and HRT changes on anaerobic fermentation process for biogas production. The parameters that were analyzed, as well as the corresponding applied standard analytical method or widely accepted protocol, are presented in Table 2.

**Table 2.** Analyzed parameter and applied analytical method used in the study.

| Parameter | Applied analytical method | Parameter | Applied analytical method |
| --- | --- | --- | --- |
| COD | APHA 5220-C | *Crude Protein | ISO 5983-2:2009 |
| COD soluble | APHA 5220-D | *Crude Fat | ISO 6492:1999 |
| *Total Solids at 103–105 °C | APHA 2540-B | *Crude Fiber | ISO 6865:2000 |
| Volatile solids (VS) at 550 °C | APHA 2540-E | Nitrogen |  |
| Total Suspended Solids (TSS) at 103–105°C | APHA 2540-D | Free Extracts (NFE) |  |
| Volatile Suspended Solids (VSS) | APHA 2540-E | Calcium (Ca) | ICP-MS using an Agilent 7850 ICP-MS with ORS4 collision cell, and following the digestion procedures outlined in ISO 17294-2:2016 (ISO, 2016) and APHA 3125 |
| Total Nitrogen (N) | APHA 4500-Norg B | Sulfur (S) |  |
| Ammoni Nitrogen (N-NH<sub>4</sub>) | APHA 4500-NH<sub>3</sub> B & C | Magnesium (Mg) |  |
| Total Phosphorus (P) | APHA 4500-P B & E | Lead (Pb) |  |
| Orthophosphates (PO<sub>4</sub>) | APHA 4500-P E | Cadmium (Cd) |  |
| pH | APHA 4500-H+ B | Chromium (Cr) |  |
| Conductivity at 25°C | APHA 2510 B | Mercury (Hg) |  |
| Salinity | APHA 2520 B | Nickel (Ni) |  |
| TAC- Total Alkalinity of Carbonates |  | Internal method |  |
| FOS-Volatile fatty acids |  | Sodium (Na) |  |
| FOS/TAC ratio |  | Potassium (K) |  |
| Acetic acid |  | Arsenic (As) |  |
| Acetic acid equivalent |  | Copper (Cu) |  |
| Propionic acid |  | Zinc (Zn) |  |
| Isobutyric acid |  | Detection of Salmonella spp | ISO 6579-1:2017 |
| Butyric acid |  | Escherichia Coli | ISO CEN/TR 16193 |
| Isovaleric acid |  | Enterococcus faecalis | ISO 7899-2/2000 |
| n-Heptanoic acid |  | Microbiome analysis |  |
| Valeric acid |  | Biochemical methane potential (BMP) | Internal method |
| Isocaproic acid |  |  |  |
| Hexanoic acid | APHA 5220-C |  |  |

\* Data used for the determination of theoretical biogas yield of organic matter (OM) by determining the contents of fat, protein, and fibrous substances per dry matter (DM) mass, and by following Baserga (1998) equation (1).

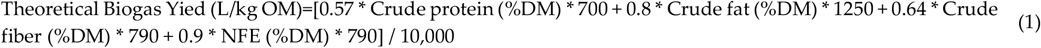

Additionally, the biochemical methane potential (BMP) of SB1 and SB2, i.e. the experimental biogas yield that is defined as the volume of methane produced per unit amount of organic substrate material added to a reactor, was determined using a Bioprocess Control AMPTS III system and by applying the measurement protocols established by the manufacturer. In more detail, 500 mL Duran Schott bottles, with a working volume of 400 mL and a headspace of 100 mL are immersed in a thermostatic water bath and periodically agitated via mechanical stirrer. The headspaces of the bottles are equipped with gas outlets that are connected to a gas volume measuring apparatus following the water displacement method. Blank samples, i.e. samples containing only inoculum, were analyzed along substrate samples, while all headspaces were purged with nitrogen gas for 2 min prior analysis initiation. All samples remained at a constant temperature of 40oC for a hydraulic retention time (HRT) of 30 days. The quantity of substrate to be added in each bottle was determined based on the following equation (2).

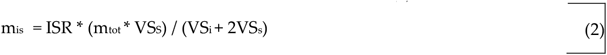

Where: m_is_ is the mass of inoculum in grams; ISR is the VS_i_ to VS_s_ ratio coefficient, set to a value of 2; mtot is the total mass that is placed in 400ml bottle working volume; VS_i_ is the % fraction of inoculum volatile solids; VS_s_ is the % fraction of substrate volatile solids.

The BMP is then calculated based on the following equation (3).

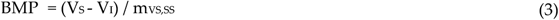

Where: V_s_ is the accumulated volume of biomethane from the reactor containing the sample (substrate and inoculum); V_i_ is the volume of biomethane produced by the inoculum quantity present in the samples; m_VS,SS_ is the mass of substrate organic material contained in the 400ml bottle working volume.

The biogas produced during the BMP measurement is collected in gas able bags and transferred for analysis using a gas chromatographer, where CH_4_, CO_2_, CO, H_2_S, NH_3_, H_2_ are quantified.

In the CSTR1 and CSTR2 experimental configurations, the gaseous products are channeled to a biogas analyzer, whereas through the use of a triode valve, biogas is collected in 2-litre gas able bags for further analysis and control of the biogas analyzer’s outputs.

Both CSTR1 and CSTR2 experimental configurations are equipped with volumetric flow meters that follow the fluid displacement method. Prior CSTR1 and CSTR2 initiation, additional tests to evaluate the accuracy of volumetric flow measurement units and to identify potential leakages from the experimental configurations were performed.

The gaseous products of CSTR1 and CSTR2 are channeled to a biogas analyzer, whereas using a triode valve, biogas could be collected in 2-litre gas able bags for further analysis and control of the biogas analyzer’s outputs (allowed discrepancies in measured concentrations <5%).

The liquid samples from CSTR1 and CSTR2 were collected from the discharge of the bioreactors (discharge pump), which occur simultaneously to their feeding (feeding pump), and their analysis was immediately routed and performed at the accredited according to ISO 17025 analytical laboratory of QLAB, located at the same facilities where both experimental CSTRs are installed and operated.

The obtained measurement data were statistically interpreted using SPSS20 software. Shapiro-Willlk test was used to check data distribution, and depending on whether the data are normally distributed or not, the T-tests, ANOVA and Pearson Correlation Coefficient Test, or Mann-Whitney U Test, Kruskal-Wallis Test and Spearman’s Rank Correlation Coefficient Test are used respectively.

For DNA Extraction and 16S rRNA Gene Amplicon Sequencing, genomic DNA was extracted from the CSTR samples with the DNeasy PowerSoil Pro Kit (QIAGEN, Hilden, Germany) according to the manufacturer’s instructions. The quantity and quality of the extracted DNA were then estimated using a V-630 Spectrophotometer (JASCO, Inc., Tokyo, Japan). Library preparation was performed following the standard guidelines of the 16S Metagenomic Sequencing Library Preparation protocol (IlluminaTM, Inc., San Diego, CA, USA). In brief, DNA was amplified using the HotStarTaq® Master Mix Kit (QIAGEN, Hilden, Germany) with the addition of the 341f/805r primer pair, which targets the bacterial and archaeal V3–V4 hypervariable regions of the 16S rRNA gene (341f 5′-CCTAC-GGGNGGCWGCAG-3′, 805r 5′-GACTACHVGGTATCTAATCC-3′). The PCR mixture (25 μL) contained 12.5 μL of HotStarTaq Master Mix, 5 μL of each primer and 2.5 μL of DNA (5 ng/μL). Thermal cycling conditions included an initial 3 min step at 95 °C, followed by 25 cycles of denaturation at 95 °C for 30 s, annealing at 55 °C for 30 s and elongation at 72°C for 30 s and a final extension step at 72 °C for 5 min. PCR amplicons were cleaned up by AMPure XP beads (Beckman Coulter, Brea, CA, USA) to remove unbound primers and primer dimers. Next, dual indices and Illumina sequencing adaptors were attached with an index PCR using the Nextera XT Index Kit (Illumina Inc., San Diego, CA, USA). The PCR reaction mixture (50 μL) comprised 25 μL of HotStarTaq Master Mix, 5 μL of each index, 10 μL of PCR Grade Water and 5 μL of the previous PCR product and the cycling conditions remained the same as that of the first PCR reaction except that the number of iterative cycles was reduced to 8. Afterward, indexed PCR amplicons were cleaned up using the AMPure XP beads (Beckman Coulter, Brea, CA, USA). The produced DNA lbraries were quantified with the Qubit™ 4 Fluorometer (Thermo Fisher Scientific, Waltham, MA, USA) and their size was verified via a 1.5% agarose gel electrophoresis. Equimolar concentrations of the libraries were then pooled together and a quantitative PCR was performed using the QIAseq Library Quant Assay Kit (QIAGEN, Hilden, Germany) for library concentration evaluation. The pooled library was subsequently spiked with 25% phiX control library (Illumina Inc., San Diego, CA, USA), denatured and diluted to a final concentration of 6 pM. Sequencing was performed on an Illumina MiSeqTM platform with the MiSeq Reagent Nano Kit version 2 (500-Cycle)/MiSeq Reagent Kit version 3 (600-Cycle) chemistry for a paired-end, 2D250-bp/2 × 300 cycle run.

The primer sequences were removed and reads with a low-quality score (average score, <20) were filtered out using the FASTQ toolkit within BaseSpace version 2.2.0 (IlluminaTM, Inc., San Diego, CA, USA). The 16S Metagenomics application (version 1.0.1) within BaseSpace was used to perform a taxonomic classification, which uses an Illumina-curated version of the GreenGenes taxonomic database and the RDP naive Bayes taxonomic classification algorithm with an accuracy of >98.2% at the species level (Wang et al. 2005).

## 3. Results

The CSTR1 and CSTR2 monitoring data from the 81-days experimental run were used to assess the effect of anaerobic bioreactor feeding patterns, both in terms of quantity and quality. Their statistical interpretation, along with the obtained data from microbiome analyses, offers valuable insights regarding the complex mechanisms that affect quality and quantity of biogas.

### 3.1. Evaluation of CSTR1 and CSTR2

The evaluation of statistical differences between CSTR1 and CSTR2 revealed that all the distribution of the measured physicochemical and biological parameters is the same across the value groups corresponding to CSTR1 and CSTR2 (Kruskal-Wallis Test at Significance level 0.05). Moreover, the most critical parameters that presented significant temporal variation (ANOVA test at Significance level 0.05) were the pH, TAC, NH4-N and Total Nitrogen, which along with the elative stable values of volatile fatty acids (FOS) resulted in a declining FOS/TAC ratio (Figure X).

**Figure 2.**
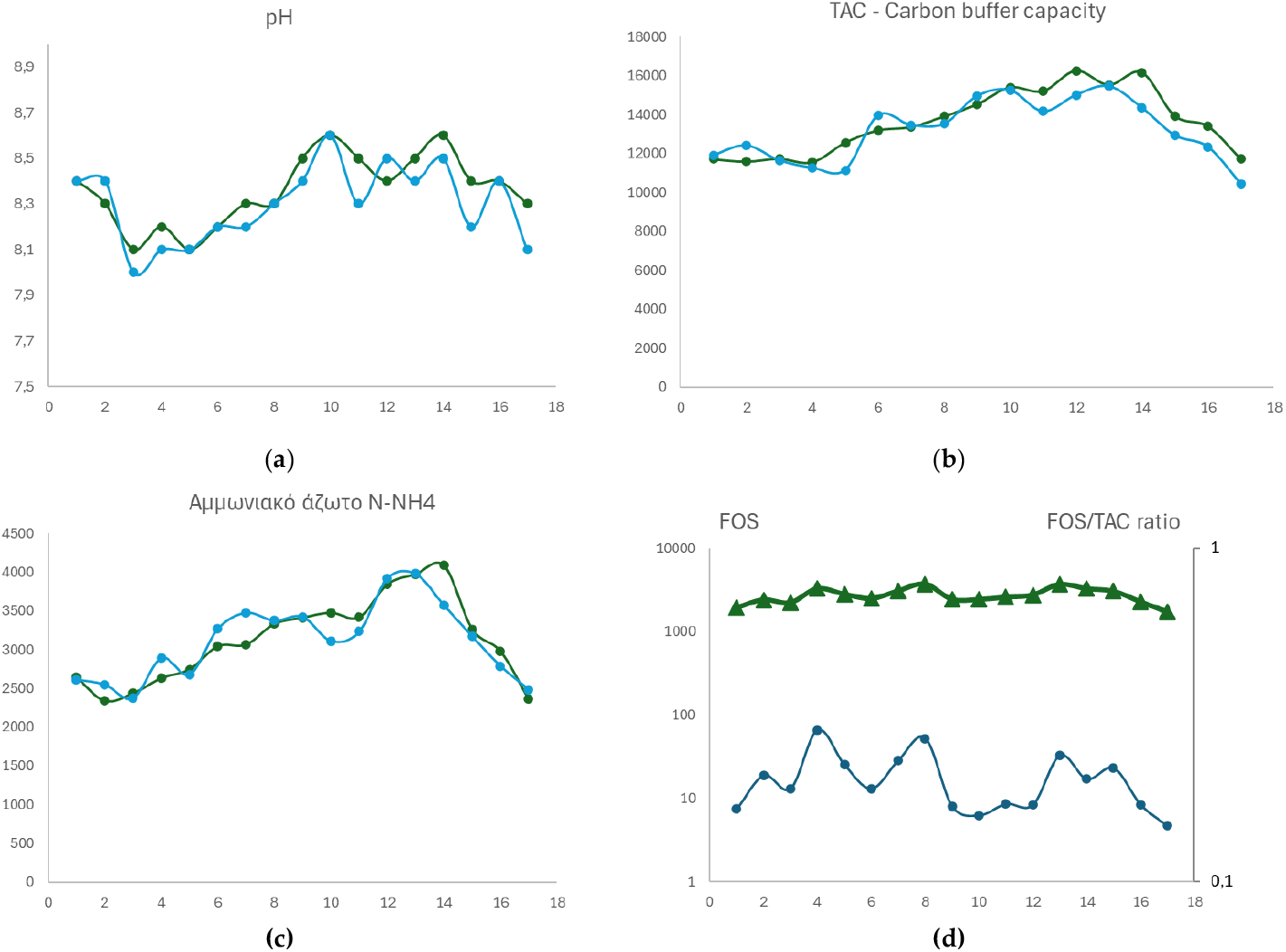
Critical monitored parameters at CSTR1 and CSTR2 that exhibit significant temporal variation. (**a**) pH; (**b**) TAC; (NH_4_-N); (**d**) FOS and FOS/TAC ratio.

According to the obtained experimental results, the parameter of TAC enhances CH_4_ content of the produced biogas, as it is evident from the Spearman’s Rank Correlation Coefficient Test, conducted at Significance level 0.01, which revealed a very strong and significant positive correlation between the measured total alkalinity of carbonates (carbon buffer capacity) and the measured CH_4_ content of produced biogas (p = 0.827, Sig.= 0.001). Furthermore, the only parameter that presented a significant correlation with TAC was the concentration of isocaproic acid, which exhibited moderately strong, statistically significant positive correlation (p = 0.647, Sig.= 0.001).

### 3.2. Biogas production in CSTR1 and CSTR2

As graphically illustrated in Figure X, during the first 24 days of the experiment (Period 1), when HRT was maintained at 20 days and SB1 was the feed of the anaerobic bioreactors, the average biogas production per unit of VS in CSTR1 and CSTR2 was 393 ml/g and 358 ml/g respectively, while the corresponding CH_4_ content was 51.4% and 53.9% respectively. While keeping an HRT of 20 days, and after the introduction of the new feed (SB2) in CSTR1 and CSTR2 (period 2), the average biogas production per unit of VS in was 364 ml/g and 349 ml/g respectively, having an average CH_4_ content of 50.3% and 52.3% respectively. The test of variation between the two first periods revealed no significant statistical difference between the data sets, thus there was an apparent acclimatization of biomass to the new feedstock. On the contrary, the data sets of the third period (period 3) of the experiment, when CSTR1 and CSTR2 were kept fed with SB2 and HRT was regulated at 30 days, revealed significant statistical variations compared to both period 1 and period 2 data sets for biogas production and methane content. The average biogas production per unit of VS in during the third period was 424 ml/g and 447 ml/g respectively, having an average CH_4_ content of 57,7.3% and 56.3% respectively.

**Figure 2.**
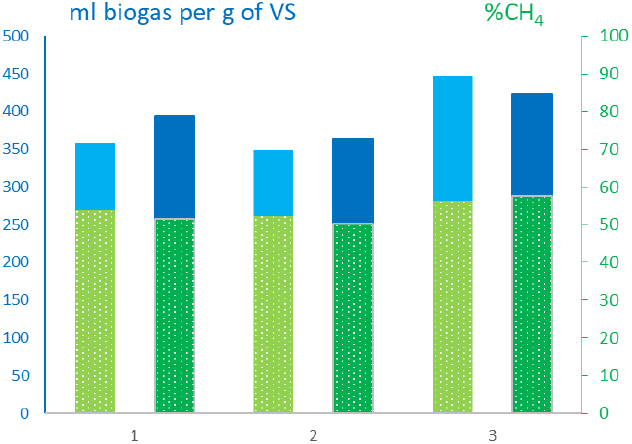
Biogas production in CSTR1 and CSTR2 during the three distinguished periods of experimental runs.

### 3.3. Microbiome of CSTR1 and CSTR2

The evaluation of statistical differences between the microbiome of CSTR1 and CSTR2 revealed that all the distribution of the measured physicochemical and biological parameters is the same across the value groups (Kruskal-Wallis Test at Significance level 0.05), thus no significant differences between the anaerobic reactors. However, the data sets from microbiome analysis presented significant temporal variation (ANOVA test at Significance level 0.05) in taxa and microbial species (Figure X). Moreover, the test of variation between the microbiome of the three periods of experiment, where different HRT or feedstock was applied, revealed significant statistical differences between all three data sets. These variations in microbiome are graphically illustrated in Figure X.

**Figure 2.**
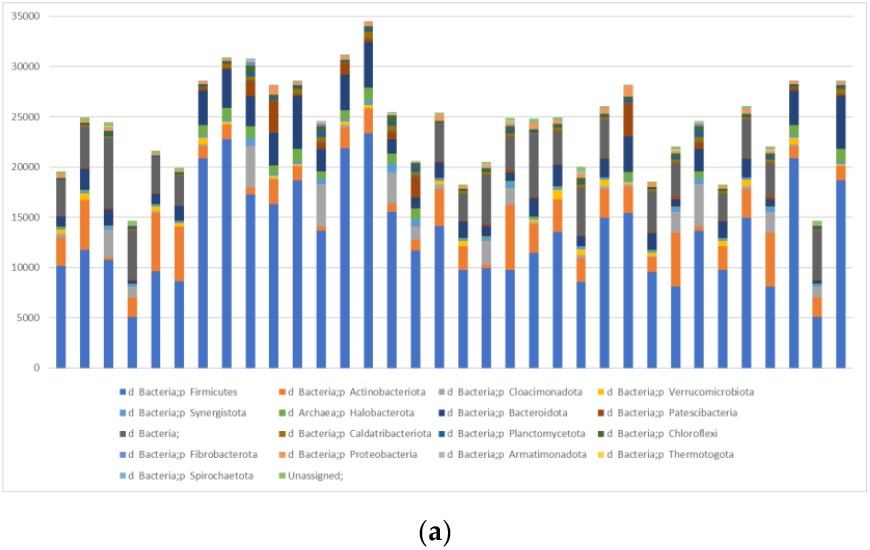
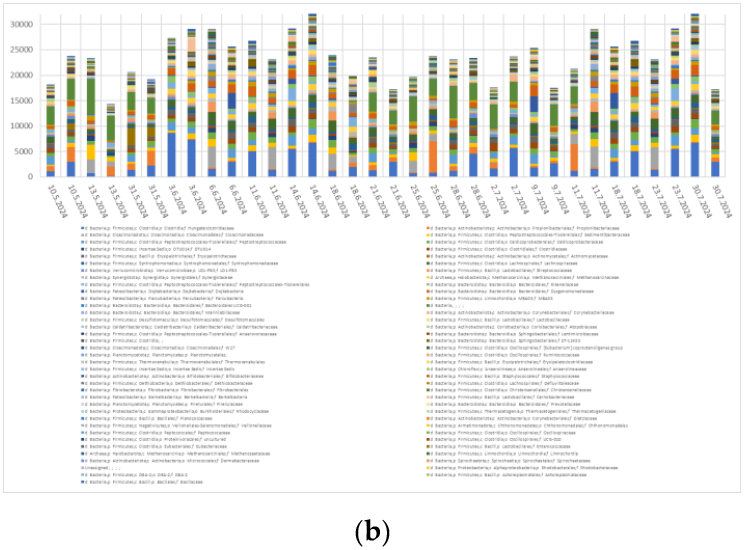
Bioinformatics of CSTR1 and CSTR2. (a) Higher Rank; (b) Lower Rank.

## 5. Conclusions

CSTR1 and CSTR2 presented similar characteristics regarding the aspects of their physicochemical characteristics, biogas production and composition, and microbiome. In both CSTR1 and CSTR2, the introduction of SB2, which has a higher content of ammonia-rich poultry manure, as well as of protein-rich food waste, resulted in increased pH and TAC values. The effect of TAC increase can be indicated in the declining FOS/TAC ratio that could eventually fall outside the desired range of values if no adjustments are made.

In full scale biogas applications, a change of feeding to overcome the FOS/TAC decline is not always achievable. The results of the study suggest the increase of HRT is an alternative measure that can stabilize the decline of FOS/TAC ratio.

The apparent loss of biogas production due to lower feeding rate at increasing HRT is compensated to a significant degree by the higher biogas production and biomethane content per unit of volatile solids that is observed at increasing HRT. In the case of CSTR1 and CSTR2, this increase was 21-30% in terms of biomethane yield.

A significant finding of the study is that the increase of Archaea Halobacterota, which are associated with the methanation step of biogas production process, does not increase biomethane content of biogas. The significant increase of their presence during the second period of experiment did not result to an increase of biogas quality. Their presence is important to finalize the process, but other taxa are seemingly foster the anaerobic process to increase the quality of the produced biogas.

In the case of CSTR1 and CSTR2, the taxa that had a statistically significant effect on biogas quality, i.e. biomethane content, are d Bacteria involved in all biochemical steps prior methanogenesis.

## Disclaimer/Publisher’s Note

The statements, opinions and data contained in all publications are solely those of the individual author(s) and contributor(s) and not of MDPI and/or the editor(s). MDPI and/or the editor(s) disclaim responsibility for any injury to people or property resulting from any ideas, methods, instructions or products referred to in the content.

